# Genomic analysis of human polymorphisms affecting drug-protein interactions

**DOI:** 10.1101/119933

**Authors:** Oriol Pich i Rosello, Anna V. Vlasova, Polina A. Shichkova, Yuri Markov, Peter K. Vlasov, Fyodor A. Kondrashov

## Abstract

Human genetic variability is thought to account for a substantial fraction of individual biochemical characteristics – in biomedical sense, of individual drug response. However, only a handful of human genetic variants have been linked to medication outcomes. Here, we combine data on drug-protein interactions and human genome sequences to assess the impact of human variation on their binding affinity. Using data from the complexes of FDA-drugs and drug-like compounds, we predict SNPs substantially affecting the protein-ligand binding affinities. We estimate that an average individual carries ~6 SNPs affecting ~5 different FDA-approved drugs from among all of the approved compounds. SNPs affecting drug-protein binding affinity have low frequency in the population indicating that the genetic component for many ADEs may be highly personalized with each individual carrying a unique set of relevant SNPs. The reduction of ADEs, therefore, may primarily rely on the application of computational genome analysis in the clinic rather than the experimental study of common SNPs.

---

Adverse drug effects (ADEs), instances when medication causes an unintended adverse response, substantially contribute to morbidity, the cost and time of treatment (Rodríguez-Monguió et al. 2003; Boeker et al. 2013) often appear unpredictably (Jolivot el al. 2014; Evans et al. 2003). Pharmacogenomics approaches identified the basis of individual drug response to several drugs (Wang et al. 2011; Relling et al. 2015), including those used in chemotherapy (Patel et al. 2015; Hertz et al. 2015). Most approaches focus on the metabolic aspect of individual drug response, leading to different dosage requirements depending on the genetic variation that affects the rate of drug metabolism (Relling et al. 2015; Hertz et al. 2015; Zhou et al. 2015). However, our understanding of the contribution of genetics to the causes and the incidence of individual drug response remains fragmented (Wang et al. 2011; Relling et al. 2015; Mooney 2015). Here we utilize a structural approach to the study of occurrence of genetic variation with probable effect on individual drug response in the human population. The molecular mechanism of action for therapeutic small molecular compounds is directly related to the strength of interaction of the drug with its intended target, typically a protein (Hopkins et al. 2011; Swinney et al. 2004). Thus, to assay the prevalence of human variability potentially affecting drug response we combined genome-wide data on human single nucleotide polymorphisms (SNPs) with structural data on drug-protein complexes.

We obtained data on human SNPs from the 1000 genome project (Auton et al. 2015), which included polymorphism data from 2504 individuals. We then selected all small molecular compound-protein structural complexes from PDB (Berman et al. 2000) in which the compound was identical or highly similar (see Methods) to an FDA-approved or an FDA experimental drug. Using the drug-protein structural complexes we identified the binding sites of the protein in direct interaction with the drug, whereby the closest heavy atom of the amino acid residue was <6 Å away from the heavy atom of the drug. We then mapped the structural data to the human genome obtaining data on drug binding sites carrying polymorphisms among the 2504 sequenced individuals in our data.

## Results

Drug-protein complexes were available for ~16% of FDA-approved drugs (296/1826), and for 170 of them (~9% of the total) our approach identified a SNP in the drug binding interface. The fraction of available drug-protein complexes was substantially higher for FDA experimental drugs, ~45% (2204/4925), with ~20% of these drugs (958/4925) having a SNP in the binding interface in our data. Many of the SNPs in drug-binding sites may not influence the drug-protein interaction and, therefore, may not to contribute to ADEs or individual drug reactions.

Therefore, our idea was to use a docking approach (Morris et al. 2009) to calculate the difference in the free energy of the drug and wild-type protein interaction and that of the drug with the protein sequencing incorporating the amino acid polymorphism (Table 1).

**Table 1:** The prevalence of SNPs in proteins in complex with FDA approved and experimental compounds.

|  | **Total** | **Average/person** |
| --- | --- | --- |
| **Number of FDA drugs (in complex with with PDB structure, with a SNP in at least one binding site)** | FDA-approved: 1826<br>(296, 170)<br>FDA experimental:<br>4925 (2204, 958)<br>FDA all: 6751 (2500,<br>1128) | NA |
| **Number of SNPs at binding sites (homozygous / heterozygous)** | FDA-approved: 554<br>(26/552)<br>FDA experimental:<br>2766 (186/2757)<br>FDA all: 3052<br>(199/3042) | FDA-approved: 3.33<br>(0.59/2.74)<br>FDA experimental: 29.87<br>(5.11/24.76)<br>FDA all:<br>31.47 (5.5/26) |
| **Number of SNPs probably affecting drug-protein binding affinity (homozygous / heterozygous)** | FDA-approved: 53 (0/53)<br>FDA-experimental: 390 (15/390)<br>FDA all: 425 (15/425) | FDA-approved: 0.03 (0/0.03)<br>FDA-experimental: 3.1 (0.61/2.49)<br>FDA all: 3.12 (0.61/2.51) |
| **Number of SNPs possibly affecting drug-protein binding affinity (homozygous / heterozygous)** | FDA-approved: 192 (5/191)<br>FDA-experimental: 835 (41/833)<br>FDA all: 956 (45/953) | FDA-approved: 0.81 (0.1/0.71)<br>FDA-experimental: 4.45 (0.46/3.9)<br>FDA all: 5 (0.55/4.45) |
| **Number of drugs probably (possibly) affected by a SNP** | FDA-approved: 30 (75)<br>FDA experimental: 165 (357)<br>FDA all: 195 (432) | FDA-approved: 0.04 (0.79)<br>FDA-experimental : 3 (4.9)<br>FDA-all : 3.04 (5.69) |

The difference in the free energy, ΔΔG, represents the difference in the strength of binding of the drug to the protein, which we then used as a measure of the impact of the SNP on the function of the protein target. A vast majority of the SNPs in drug-protein binding sites were predicted not to affect binding affinity. However, many more SNPs were predicted to decrease the strength of interaction rather than increase it (Figure 1).

**Figure 1.**
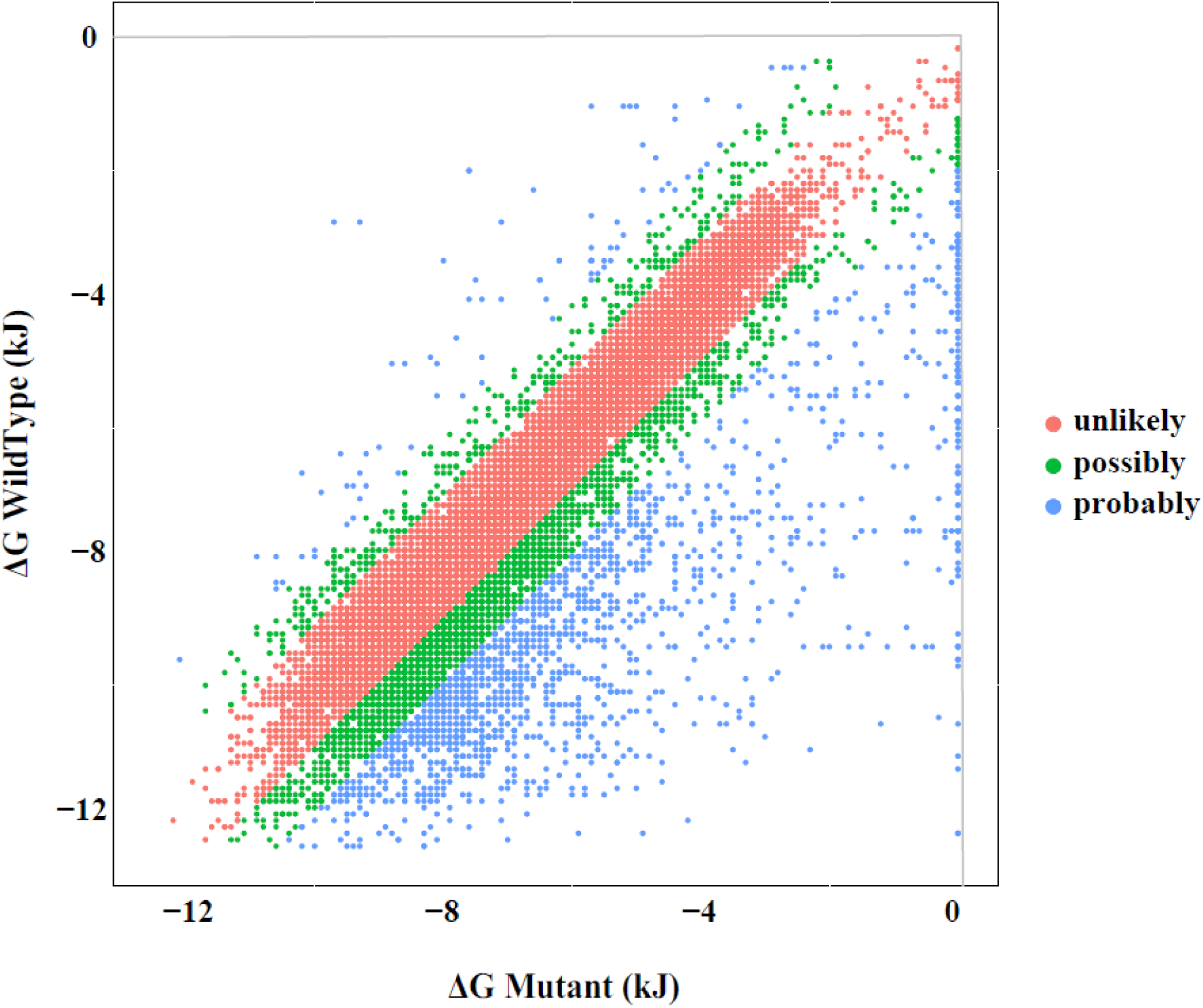
The effect of SNPs in protein binding sites on the binding strength of drug ligands. X-axis represents the **Δ**G of the drug-protein complex with the major allele in the polymorphic site. The Y-axis represents the **Δ**G of the same complex with the minor allele at the same site.

Existing docking methods do not always provide an accurate estimate of the free energy of the interaction (Sousa et al. 2013). Therefore, we validated our method using two separate datasets. First, we used FDA guidelines on genotype (table of pharmacogenomic biomarkers in drug labeling) to obtain a list of compounds for which the FDA recommends genetic testing (FDA web, 2016). From the list we selected 5 SNPs in two Cytochrome P450 proteins and in EGFR that are known to cause an individual response to 7 different FDA-approved compounds. For other examples cited by the FDA either the structure of the drug-protein complex was not available, the described SNPs did not occur in the drug-protein binding interface, or the FDA recommended screening for a haplotype containing several polymorphisms rather than a specific SNP. We found that all of the 7 interactions, from which contribution to ADEs is well documented (Marez et al. 1997; Maekawa et al. 2006; Dai et al. 2014), the minor allele SNPs caused a > 0.6 change in ΔΔG, with the average effect being around 1.5 ΔΔG (Table 2), confirming that our method identifies SNPs known to affect individual drug response as having an effect on ΔΔG.

**Table 2.** Estimated ΔΔG of polymorphisms recommended to be genotyped by the FDA in certain drug prescriptions.

| Chain | Drug_ID | Complex | Target | Emut | Enat | DD | Description PDB | Drug Name |
| --- | --- | --- | --- | --- | --- | --- | --- | --- |
| 1XKK_A | DB01259 | 1XKK_A_L858R_DB01259 | EGFR | -8.9 | -11.2 | 2,3 | A unique structure for epidermal growth factor receptor bound to Lapatinib | Lapatinib |
| 4WNU_C | DB00908 | 4WNU_C_I297L_DB00908 | CYP2D6 | -8.1 | -9.7 | 1,6 | Human Cytochrome P450 2D6 Quinidine Complex | Quinidine |
| 4WNU_C | DB00908 | 4WNU_C_R296C_DB00908 | CYP2D6 | -8.5 | -9.7 | 1,2 | Human Cytochrome P450 2D6 Quinidine Complex | Quinidine |
| 4WNW_B | DB00679 | 4WNW_B_I297L_DB00679 | CYP2D6 | -8.4 | -9.2 | 0,8 | Human Cytochrome P450 2D6 Thioridazine Complex | Thioridazine |
| 4WNW_B | DB00679 | 4WNW_B_R296C_DB00679 | CYP2D6 | -8.6 | -9.2 | 0,6 | Human Cytochrome P450 2D6 Thioridazine Complex | Thioridazine |
| 1OG5_B | DB00682 | 1OG5_B_G98V_DB00682 | CYP2C9 | -8.8 | -10.7 | 1,9 | Crystal Structure of Human Cytochrome P450 2C9 with Bound Warfarin | Warfarin |
| 1OG5_B | DB00682 | 1OG5_B_Q214L_DB00682 | CYP2C9 | -8.9 | -10.7 | 1,8 | Crystal Structure of Human Cytochrome P450 2C9 with Bound Warfarin | Warfarin |

Second, we obtained a list of mutations observed in human cancers by The Cancer Genome Atlas (TCGA) consortium (Weinstein et al. 2013). We reasoned that recurrent mutations in different human cancers, often predicted to be cancer driver mutations (Weinstein et al. 2013), are likely to have a stronger than average effect on the free energy of the binding of the protein with anticancer drugs or naturally occurring small molecular compounds. Alternatively, mutations that are found only occasionally in human cancers, likely cancer passenger mutations (Weinstein et al. 2013), should be less likely to affect the binding affinity of human proteins with small ligands. We found that mutations observed in >10 human cancers have a 2-fold higher impact on the binding of proteins with small molecular compounds than mutations found once or twice in the TCGA data (Figure 2), confirming that our method can distinguish between the binding strength of cancer driver and passenger mutations.

**Figure 2.**
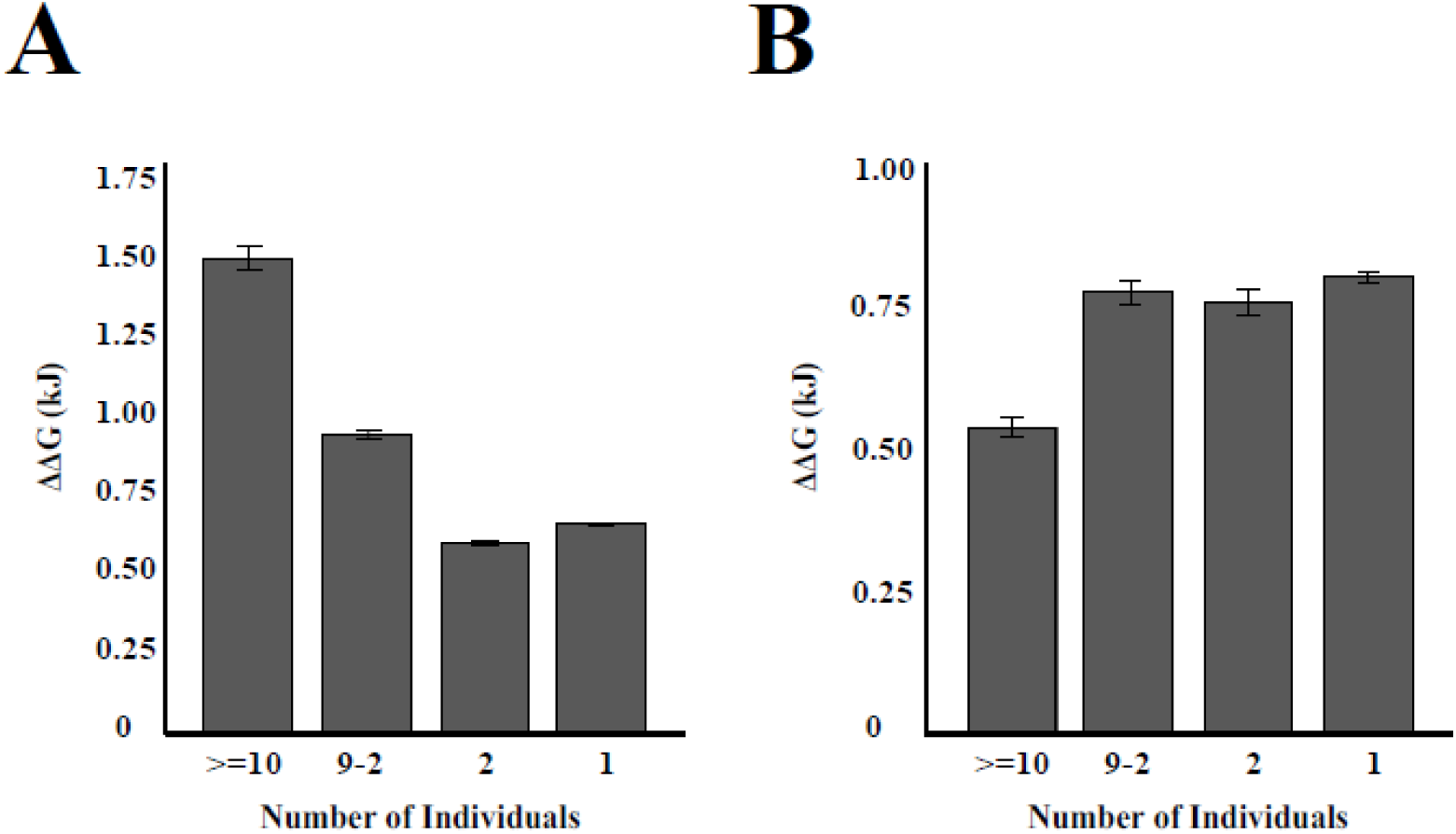
The average ΔΔG as a function of variant frequency. (a) The average effect of a cancer mutation on ΔΔG as a function of the number of cancers in which it was found. (b) The average effect of a SNP from the human population on ΔΔG as a function of the number of individuals in which the SNP was found. Error bars represent s.e.m.

Taken together, these data indicate that our approach identifies SNPs that have an effect on the known drug-protein complexes. A change in ΔΔG |2.0| kJ typically corresponds to a change in binding affinity with a high probability (Trott et al. 2010), therefore, we classified SNPs with such an effect as probably affecting the interaction of the drug with the target protein. SNPs causing a weaker change in free energy, |1.0| ΔΔG < |2.0|, were classified as possibly influencing the drug-protein interaction. All other SNPs, those predicted to have a < 1.0 effect on ΔΔG, were labeled as not likely to influence the drug-protein interaction. Our approach appears conservative, as even small changes in binding affinity, smaller than our cutoff values, are known to have an effect on the functional impact of the interaction of proteins and small molecular compounds (Hopkins et al. 2014).

Among the 2504 available human genomes, our approach identified 53 SNPs that probably affect a drug-protein interaction with 30 FDA-approved drugs and 192 SNPs that possibly have this effect on 75 FDA-approved drugs (Figure 3, Table 3).

**Figure 3.**
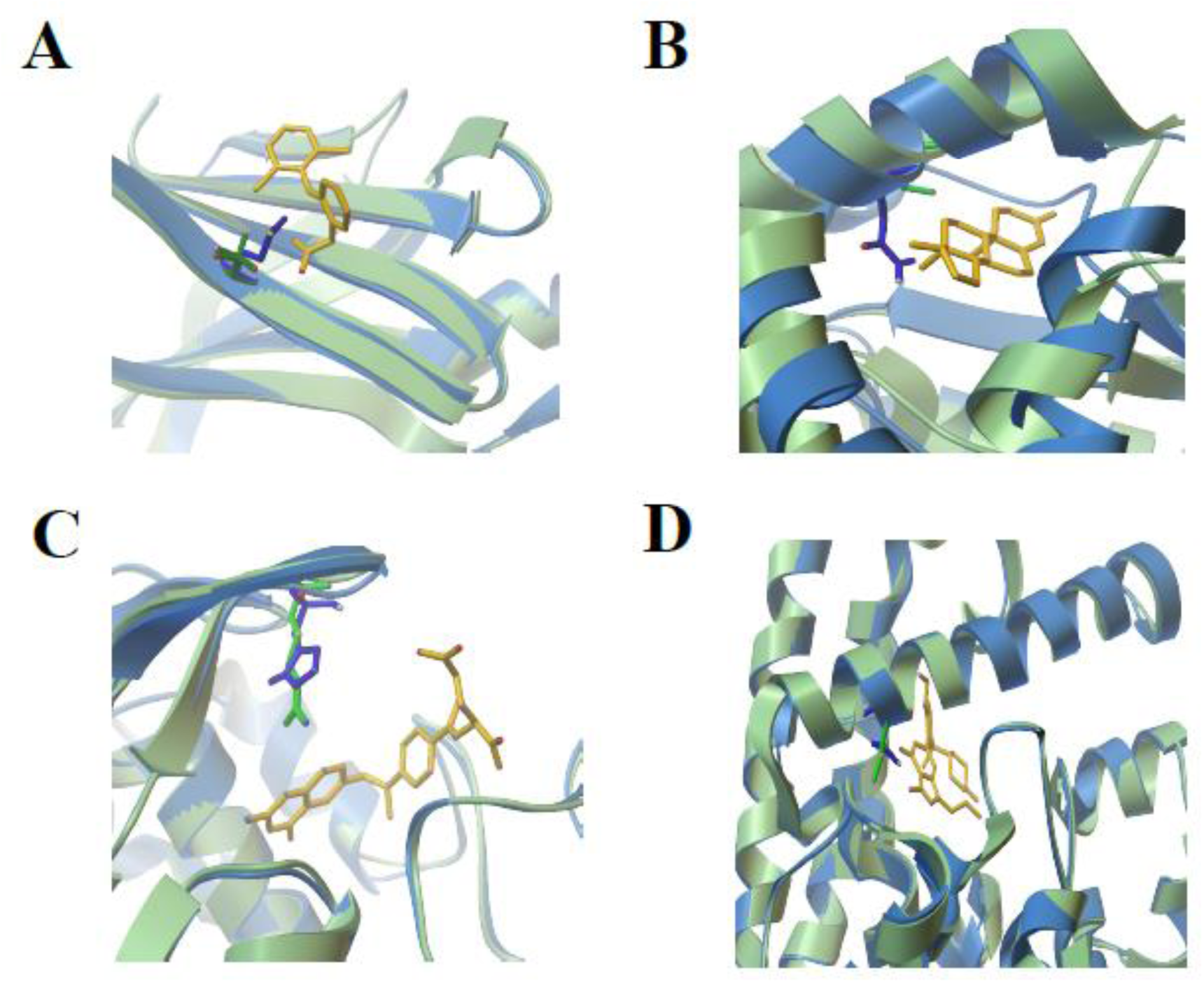
Structures of selected SNPs with a predicted effect on FDA-approved drugs. Structure with the major allele, corresponding to wild-type sequence, is shown in green, the minor allele in blue. (**a**) Diclofenac, (**b**) Testosterone, (**c**) Methotrexate and (**d**) Sildenafil are shown in yellow.

**Table 3.** Selected polymorphisms with a predicted effect on FDA-approved drugs shown in Figure 3.

| | PDB ID | Drug | Protein | Substitution | $\Delta\Delta G$ |
| --- | --- | --- | --- | --- | --- |
| A | 1DVX | Diclofenac | Transretinin | T119M | 2.2 |
| B | 1JTV | Testosterone | 17beta-HSD1 | S222N | 5.2 |
| C | 4KN0 | Methotrexate | FOLR2 | R119H | 3 |
| D | 3JWQ | Sildenafil | PDE5 | I778T | 1.8 |

The number of SNPs predicted to have an effect on FDA experimental drugs was higher, with 390 SNPs probably affecting 165 FDA experimental drugs and 835 SNPs possibly having affecting this effect on 357 FDA experimental drugs (Table 1). An average individual carries ~1 SNPs probably and possibly affecting an FDA-approved drug interaction and ~8 such SNPs affecting FDA experimental drugs of all the drugs in complex with a protein in our dataset (Table 1, Figure 4).

**Figure 4.**
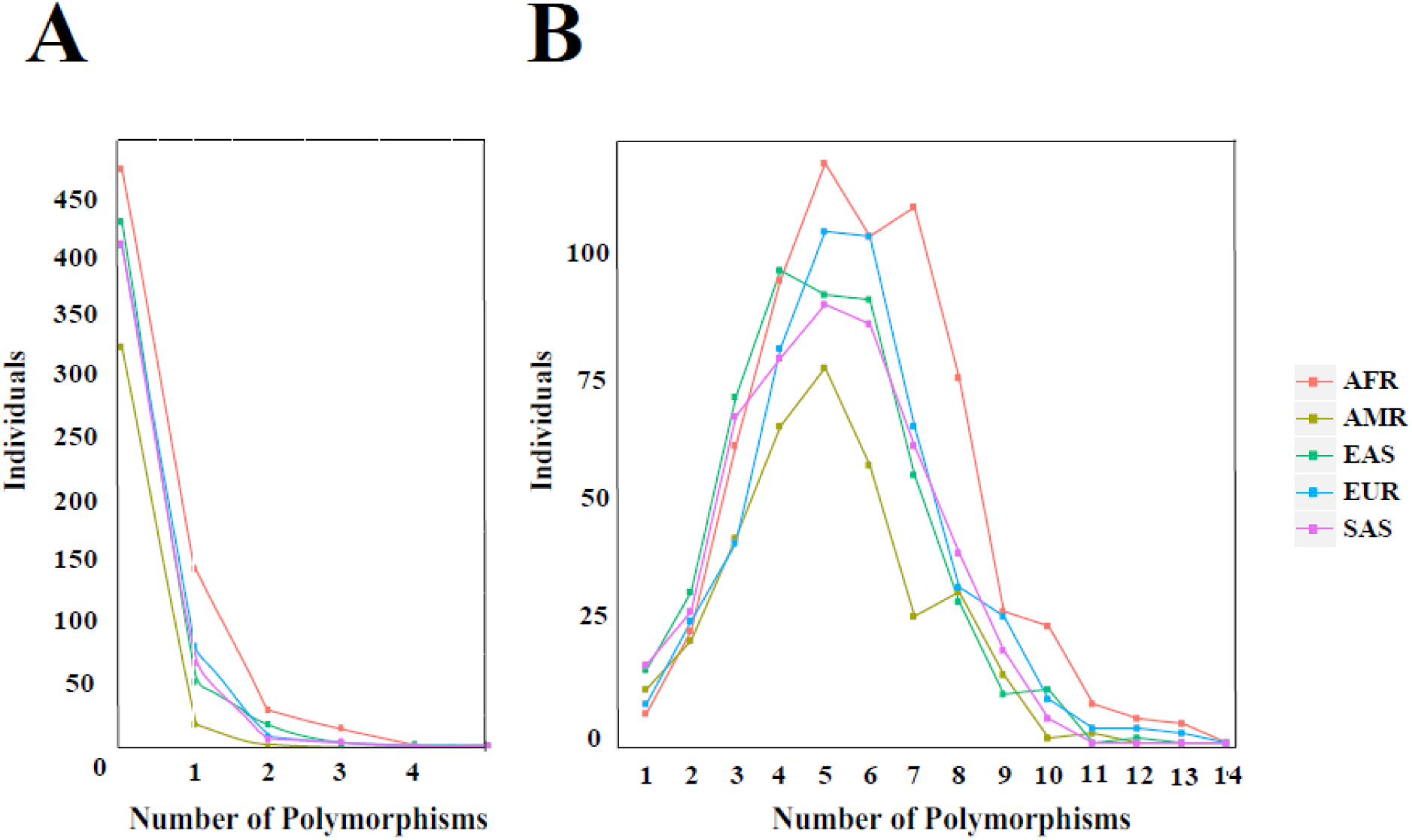
Distribution of the frequency of SNPs possibly or probably affecting a drug-protein binding affinity in human subpopulations. The distribution for (**A**) FDA-approved and (**B**) FDA experimental drugs are shown. Our data included 661 African (AFR), 347 American (AMR), 489 South Asian (SAS), 503 European (EUR) and 504 East Asian (EAS) genomes.

A majority of individuals in our data does not carry any SNPs with an effect on binding of FDA-approved compound and only 7 out of 2504 carried 5 SNPs (Figure 4A). A direct extrapolation of the distribution of SNPs with an effect to all FDA-approved drugs is complicated by the availability of the drug-protein complex for only ~15% of such drugs. However, since the average density of SNPs was the same for genes coding for proteins crystallized in complex an FDA-approved and FDA experimental drugs (4/296, 30/2204, Fisher’s exact test, p=0.2), and a drug-protein complex is available for almost one half of all experimental drugs, we used the observed distribution for 2204 FDA experimental drugs to extrapolate the expected distribution for the 1826 FDA-approved drugs.

We followed a bootstrapping approach, sampling 1826 out of 2204 FDA experimental drugs 100 times, obtaining an expected distribution of the number of SNPs with an effect on FDA-approved drug binding (Figure 5A) and the number of different FDA-approved drugs an individual may have an individual reaction to (Figure 5B).

**Figure 5.**
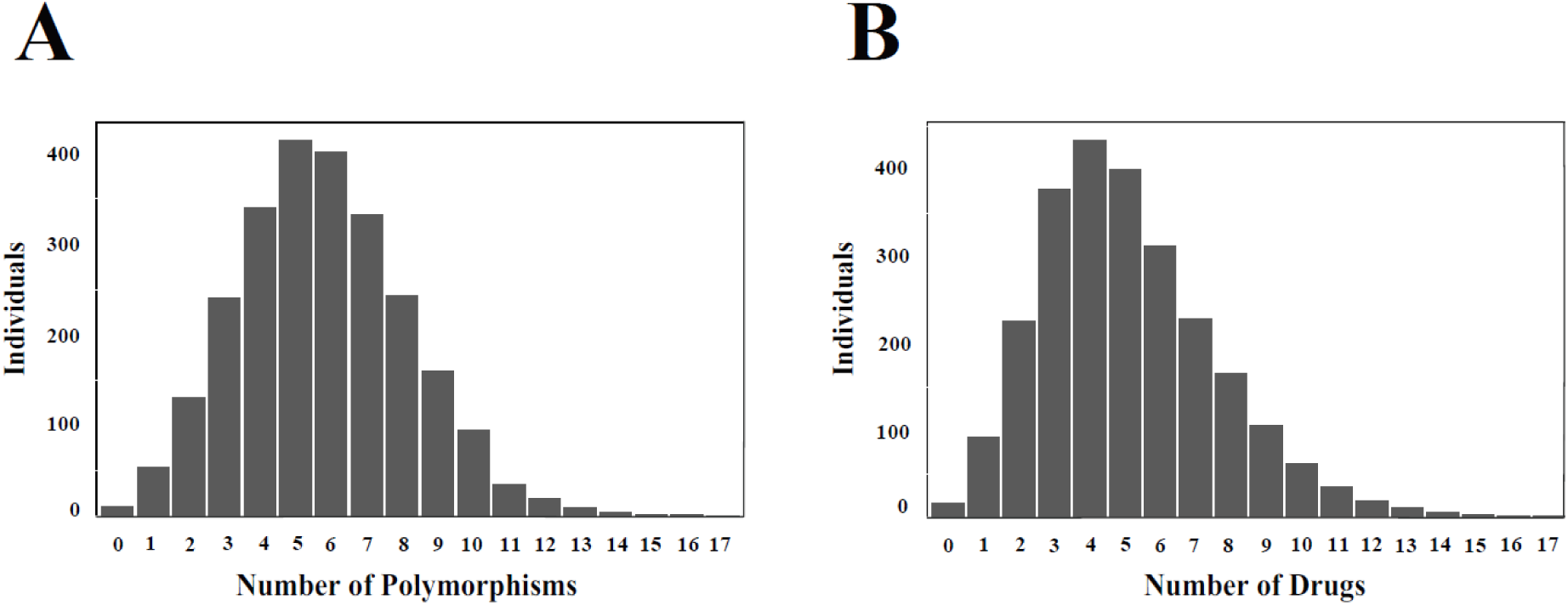
Bootstrap sampling of 2204 FDA experimental drugs in complex with a protein. The distributions of (**A**) the number of polymorphisms per genome possibly or probably affecting drug-protein binding affinity and (**B**) the number of drugs affected by such polymorphisms resulting from 1826 drugs sampled 100 times with replacement.

We predict that an average individual carries ~6 SNPs that have the potential to lead to an individual reaction to ~5 FDA-approved drugs. Interestingly, 10% of the population is expected to carry twice as many SNPs modifying an interaction with twice as many FDA-approved drugs.

SNPs with high frequency on average have a much lower effect on ΔΔG (Figure 2B). Conversely, the average frequency of SNPs probably, possibly and unlikely to have an effect was 8.4×10^−3^, 5.2×10^−3^ and 1.2×10^−2^ respectively, suggesting that such SNPs are selected against in the human population, possibly because such polymorphisms also affect the protein interaction with naturally occurring low molecular compounds. Individuals with African ancestry showed a higher number of such polymorphisms (Figure 4), which is consistent with the overall higher frequency of SNPs in the African population (Tishkoff et al. 2002). The low frequency of such polymorphisms suggest that most individuals will carry their own set with a substantial effect on drug-protein binding. Therefore, their identification through genome wide association studies or clinical trials may be too costly, requiring computational approaches, such as the one described in the present work, to provide a cost-effective means for the prediction of SNPs contributing to adverse drug-protein interaction.

## Methods

From 82494 protein-ligand structures available at Protein Data Bank, we extracted the structure of all low-molecular compounds (18535). All structures were converted from sdf to mol format using OpenBabel (O’Boyle et al. 2014), and were compared using molecular fingerprinting to 6751 FDA Drugs structures obtained from DrugBank (Wishart et al. 2006) through MolPrint2D (Bender et al. 2004). A threshold for Tanimoto Coefficient higher than 0.9 was set. We then selected the binding residues of the protein for each complex based on a less than 6 Angstroms proximity among the protein and the ligand atoms according the coordinates in the three dimensional space. FASTA sequence for all human proteins was retrieved from Ensembl (Cunningham et al. 2015), and a sequence comparison to those protein chains from PDB was performed using unidirectional BLAST (Altschul et al. 1990). Only one human protein, the one with higher bit-score, was allowed per PDB chain, but the same human protein could match different PDB chains. To assure that the binding structure was preserved in the human proteins, all PDB proteins and human proteins were aligned using MUSCLE (Edgar et al. 2004) and a filter, (length of alignment - gaps - mismatches)/length of alignment and length of alignment/PDBseq length > 0.9, was used.

A drug-protein complex was included in our study only if all the binding residues found in the drug-PDB chain interaction were also found in the human protein sequence and at least three amino acid residues per complex were found in the binding site. Proteins found in PDB that bind the same ligand and have the same binding site pattern were counted once. 1000 genome data phase 3 was retrieved and mapped to binding sites. Pan-cancer data was obtained from The Cancer Genome Atlas (downloaded in beginning 2015), and was analyzed using Ensembl VEP! (McLaren et al. 2010).

Δ*G* binding for all the drug-protein complexes (Δ*Gwt*) was calculated using a docking procedure in AutoDock (Morris et al. 2009). For each SNP observed at a binding site we modeled the original complex replacing the original binding residue for the polymorphism observed in the population, using Modeller (Eswar et al. 2007). Δ*G* binding was then calculated again for complex of the protein with the same ligand (Δ*Gm*). Finally, we subtracted the two energies obtaining ΔΔ*G*.

## Discussion

Some of the proteins in complex with FDA compounds are not the actual therapeutic targets. However, many of them may be clinically relevant because adverse drug reactions can happen due to a change in interaction of a non-therapeutical target (Wang et al. 2011; Relling et al. 2015) that can affect the drug metabolism or toxicity. Generally speaking, not all of binding proteins are equal to “final” proteins responsible for the pharmacological effect – but, any drug-protein interaction is important in terms of drugs ADME properties. Similarly, it is likely that for each of the drugs in the present study we considered only a fraction of the various interacting proteins, implying that the extrapolations presented here (Figures 5) are lower bound estimates. Finally, there are many important factors out of protein-coding regions - such as mutations in promoters, genome material rearrangements, and expression changes - that can affect disease-connected biochemical cascades and drug ADME and efficiency. Some of this factors my play the crucial role – but, in our project, we are focused on SNPs that have direct impact on proteins structures, functions and ligand binding.

Our data suggest that SNPs with a plausible contribution to ADE are present in most individuals; however, most of such polymorphisms are rare requiring a personalized approach to their identification. Importantly, given the low frequency of such SNPs in the population it is unlikely that the design of new drugs can entirely eliminate the possibility of ADEs in a small fraction of patients. Instead, the greater availability of genomic information for patients coupled with advancement of computational tools for the analysis of the effect of SNPs on drug binding may pave the way for decreasing the number of ADEs in the clinic. Specifically, clinicians should consider alternative medication if the patient carries a SNP in the drug-protein interface with the initial drug of choice, especially if the SNP is predicted to have a strong effect on the binding affinity.

## Acknowledgements

We thank Inna Povolotskaya for technical assistance. The work was supported by HHMI International Early Career Scientist Program (55007424), the EMBO Young Investigator Programme, MINECO and FEDER grant (BFU2012-31329 and Sev-2012-0208), Secretaria d'Universitats i Recerca del Departament d'Economia i Coneixement de la Generalitat’s AGAUR program (2014 SGR 0974), and the European Research Council under the European Union's Seventh Framework Programme (FP7/2007-2013, ERC grant agreement (335980_EinME). We acknowledge the support of the Spanish Ministry of Economy and Competitiveness, ‘Centro de Excelencia Severo Ochoa 2013-2017’ (SEV-2012-0208).

## The authors declare no financial interests

Correspondence and requests for materials should be addressed to

## References

Rodríguez-Monguió R, Otero MJ, Rovira J. Assessing the economic impact of adverse drug effects. Pharmacoeconomics. 2003; 21(9): 623–50.

Boeker EB, de Boer M, Kiewiet JJ, Lie-A-Huen L, Dijkgraaf MG, Boermeester MA. Occurrence and preventability of adverse drug events in surgical patients: a systematic review of literature. BMC Health Serv Res. 2013 Sep 28; 13: 364.

Jolivot PA, Hindlet P, Pichereau C, Fernandez C, Maury E, Guidet B, Hejblum G. A systematic review of adult admissions to ICUs related to adverse drug events. Crit Care. 2014 Nov 25; 18(6): 643.

Evans WE, McLeod HL. Pharmacogenomics‐‐drug disposition, drug targets, and side effects. N Engl J Med. 2003 Feb 6; 348(6): 538–49.

Wang L, McLeod HL, Weinshilboum RM. Genomics and drug response. N Engl J Med. 2011 Mar 24; 364(12): 1144–53.

Relling MV, Evans WE. Pharmacogenomics in the clinic. Nature. 2015 Oct 15;526(7573):343–50.

Patel JN. Cancer pharmacogenomics: implications on ethnic diversity and drug response. Pharmacogenet Genomics. 2015 May;25(5): 223–30.

Hertz DL, Rae J. Pharmacogenetics of cancer drugs. Annu Rev Med. 2015; 66: 65–81.

Zhou ZW, Chen XW, Sneed KB, Yang YX, Zhang X, He ZX, Chow K, Yang T, Duan W, Zhou SF. Clinical association between pharmacogenomics and adverse drug reactions. Drugs. 2015 Apr;75(6): 589–631.

Mooney SD. Progress towards the integration of pharmacogenomics in practice. Hum Genet. 2015 May;134(5): 459–65.

Hopkins AL, Groom CR. The druggable genome. Nat Rev Drug Discov. 2002 Sep;1(9): 727–30.

Swinney DC. Biochemical mechanisms of drug action: what does it take for success? Nat Rev Drug Discov. 2004 Sep;3(9): 801–8.

Auton, A. et al. A global reference for human genetic variation. Nature 526, 68–74 (2015).

Berman, H. M. The Protein Data Bank. Nucleic Acids Res. 28, 235–242 (2000).

Morris, G. M. et al. AutoDock4 and AutoDockTools4: Automated docking with selective receptor flexibility. J. Comput. Chem. 30, 2785–91 (2009).

Sousa SF, Ribeiro AJ, Coimbra JT, Neves RP, Martins SA, Moorthy NS, Fernandes PA, Ramos MJ. Protein-ligand docking in the new millennium‐‐a retrospective of 10 years in the field. Curr Med Chem. 2013; 20(18): 2296–314.

Marez D, Legrand M, Sabbagh N, Lo Guidice JM, Spire C, Lafitte JJ, Meyer UA, Broly F., Polymorphism of the cytochrome P450 CYP2D6 gene in a European population: characterization of 48 mutations and 53 alleles, their frequencies and evolution. Pharmacogenetics. 1997 Jun;7(3): 193–202.

Maekawa K, Fukushima-Uesaka H, Tohkin M, Hasegawa R, Kajio H, Kuzuya N, Yasuda K, Kawamoto M, Kamatani N, Suzuki K, Yanagawa T, Saito Y, Sawada J., Four novel defective alleles and comprehensive haplotype analysis of CYP2C9 in Japanese. Pharmacogenet Genomics. 2006 Jul;16(7):497–514

Dai DP, Xu RA, Hu LM, Wang SH, Geng PW, Yang JF, Yang LP, Qian JC, Wang ZS, Zhu GH, Zhang XH, Ge RS, Hu GX, Cai JP., CYP2C9 polymorphism analysis in Han Chinese populations: building the largest allele frequency database, Pharmacogenomics J. 2014 Feb;14(1):85–92. doi: 10.1038/tpj.2013.2.

Weinstein, J. N. et al. The Cancer Genome Atlas Pan-Cancer analysis project. Nat. Genet. 45, 1113–20 (2013).

Trott O, Olson AJ. AutoDock Vina: improving the speed and accuracy of docking with a new scoring function, efficient optimization, and multithreading. J Comput Chem. 2010 Jan 30; 31(2): 455–61.

Hopkins AL, Keserü GM, Leeson PD, Rees DC, Reynolds CH. The role of ligand efficiency metrics in drug discovery. Nat Rev Drug Discov. 2014 Feb;13(2): 105–21.

Tishkoff SA, Williams SM. Genetic analysis of African populations: human evolution and complex disease. Nat Rev Genet. 2002 Aug;3(8): 611–21.

O’Boyle NM, Banck M, James CA, Morley C, Vandermeersch T, Hutchison GR. Open Babel: An open chemical toolbox. J Cheminform. 2011 Oct 7; 3: 33.

Wishart DS, Knox C, Guo AC, Shrivastava S, Hassanali M, Stothard P, Chang Z, Woolsey J. DrugBank: a comprehensive resource for in silico drug discovery and exploration. Nucleic Acids Res. 2006 Jan 1;34

Bender A, Mussa HY, Glen RC, Reiling S. Similarity searching of chemical databases using atom environment descriptors (MOLPRINT 2D): evaluation of performance. J Chem Inf Comput Sci. 2004 Sep-Oct;44(5): 1708–18.

Cunningham F, et al. Ensembl 2015. Nucleic Acids Res. 2015 Jan;43(Database issue):D662–9.

Altschul, S.F., Gish, W., Miller, W., Myers, E.W. & Lipman, D.J. (1990) “Basic local alignment search tool.” J. Mol. Biol. 215:403–410

Edgar RC. MUSCLE: multiple sequence alignment with high accuracy and high throughput. Nucleic Acids Res. 2004 Mar 19; 32(5): 1792–7.

McLaren W, Pritchard B, Rios D, Chen Y, Flicek P, Cunningham F. Deriving the consequences of genomic variants with the Ensembl API and SNP Effect Predictor. Bioinformatics 26(16):2069–70(2010)

Eswar, N. et al. Comparative protein structure modeling using MODELLER. Curr. Protoc. Protein Sci. Chapter 2, Unit 2.9 (2007).

http://www.fda.gov/Drugs/ScienceResearch/ResearchAreas/Pharmacogenetics/ucm083378.htm (FDA Pharma)

